# Relic DNA is abundant in soil and obscures estimates of soil microbial diversity

**DOI:** 10.1101/043372

**Authors:** Paul Carini, Patrick J. Marsden, Jonathan W. Leff, Emily E. Morgan, Michael S. Strickland, Noah Fierer

**Affiliations:** Cooperative Institute for Research in Environmental Sciences, University of Colorado, Boulder, CO 80309; Department of Chemistry, Metropolitan State University of Denver, Denver, CO 80217; Department of Ecology and Evolutionary Biology, University of Colorado, Boulder, CO 80309; Department of Biological Sciences, Virginia Tech University, Blacksburg, VA 24061

## Abstract

It is implicitly assumed that the microbial DNA recovered from soil originates from living cells. However, because relic DNA (DNA from dead cells) can persist in soil for weeks to years, it could impact DNA-based analyses of microbial diversity. We examined a wide range of soils and found that, on average, 40% of prokaryotic and fungal DNA was derived from the relic DNA pool. Relic DNA inflated the observed prokaryotic and fungal diversity by as much as 55%, and caused misestimation of taxon abundances, including taxa integral to key ecosystem processes. These findings imply that relic DNA can obscure treatment effects, spatiotemporal patterns, and relationships between taxa and environmental conditions. Moreover, relic DNA may represent a historical record of microbes formerly living in soil.

**One Sentence Summary:** Soils can harbor substantial amounts of DNA from dead microbial cells; this ‘relic’ DNA inflates estimates of microbial diversity and obscures assessments of community structure.

## Main text

Microbes play critical roles in terrestrial biogeochemistry and the maintenance of soil fertility. Microbiologists, mycologists, biogeochemists, and ecologists now routinely use DNA-based approaches to determine the composition and diversity of soil microbial communities using molecular methods that include amplicon (marker gene) sequencing, quantitative polymerase chain reaction (qPCR), and shotgun metagenomics. These methods have advanced our understanding of terrestrial microbiology in a myriad of ways, by: *i)* revealing that thousands of unique microbial taxa can inhabit a single gram of soil (*1*-*3*); *ii)* uncovering novel soil microbial diversity (*4*, *5*); and *iii)* identifying putative functions of uncultivated taxa (*6*). As DNA sequencing costs continue to plummet, the use of molecular methods to describe soil microbial diversity aimed at answering both basic and applied research questions will become even more commonplace.

Linking the activities of microbes to soil processes first necessitates distinguishing living cells (both metabolically active and dormant) from those that are dead. Most molecular investigations of soil microbial diversity make the implicit assumption that the total pool of DNA extracted from soil is derived exclusively from living cells contributing to, or potentially contributing to, biogeochemical transformations. However, total microbial DNA can originate from both living and dead cells. Previous work has shown that extracellular DNA and DNA from dead or partially degraded organisms can persist in soils for weeks to years (reviewed in (*7*-*9*)). The longevity and size of this extracellular DNA pool is controlled by a myriad of soil factors. For example, complex physical factors such as soil mineralogy, pH and ionic strength control the sorption of DNA to the soil matrix, as well as the molecular integrity of the DNA itself (*10*, *11*). This sorbed DNA can be protected from removal by microbes that use it as a source of transformable genetic material or for nutrition (reviewed in (*12*)).

Given the potential for extracellular DNA to persist in soil, we hypothesized that DNA from dead microbes (which we term ‘relic DNA’) may obscure DNA-based estimates of the diversity and structure of soil microbial communities (*13*, *14*). Such effects should be most apparent if relic DNA is abundant and if the taxa represented in the relic DNA pool are not reflective of the taxa present as living cells. It is important to distinguish living microbes from dead microbes because ecological definitions of community-level diversity and structure are meant to encompass organisms actually alive at a site, not both the living and extinct organisms. Here, we used a propidium monoazide (PMA)-based approach (*15*, *16*) to remove relic DNA from a broad range of soil types. We report the amount of microbial DNA derived from relic DNA pools and show how relic DNA affects the observed richness and composition of microbial communities. We also identify which soil characteristics are linked to greater relic DNA effects and discuss the implications of relic DNA on molecular analyses of microbial communities in soil and other environments.

We tested the effect of relic DNA on estimation of microbial diversity by analyzing 31 soils collected from a broad range of ecosystem types across the United States, selected to encompass a wide variety of edaphic characteristics (Supplementary Table S1). Subsamples of each soil were either treated with PMA (n=5) or left untreated (n=5). PMA is a DNA intercalating molecule that is generally excluded by cells with intact membranes, but binds extracellular DNA and DNA of cells with compromised cytoplasmic membranes (*15*). After exposure to light, intercalated PMA covalently binds and permanently modifies DNA, rendering it un-amplifiable by the polymerase chain reaction (PCR) (*15*). We used quantitative PCR (qPCR) to calculate the amount of relic DNA by subtracting the number of amplifiable prokaryotic 16S rRNA gene copies or fungal internal transcribed spacer 1 (ITS) amplicon copies in untreated samples (total DNA = DNA from living cells + relic DNA) from the number of gene copies in PMA-treated samples (DNA from living cells only). Microbial communities were characterized by high-throughput sequencing of amplified rRNA gene regions (16S rRNA for prokaryotes or the ITS region for fungi) from both PMA treated and untreated soils. We compared estimates of microbial richness, overall community composition, and taxon abundances after standardizing all libraries to equivalent sequencing depths (see materials and methods in Supplementary Materials).

Relic DNA represented a large fraction of microbial DNA in many soils. Across all 31 soils, 40.7 ± 3.75% (mean ± SE; *n*=155) of amplifiable prokaryotic 16S rRNA genes were derived from the relic DNA pool (Fig. 1A). Similar patterns were observed for fungi, where 40.5 ± 4.12% (mean ± SE; *n*=155) of fungal ITS amplicons originated from the relic DNA pool (Fig. 1B). There are a number of lines of evidence that suggest these estimates of the amounts of relic DNA found in soil are conservative. First, when we experimentally added naked DNA to soil, our approach completely removed the naked DNA from most soil samples, but only reduced (did not completely remove) naked DNA added to soils which were found to contain high levels of relic DNA (Fig. S1A). Second, because not all dead cells have compromised membranes (*17*), PMA may not infiltrate all dead cells. Conversely, it is unlikely our approach removes DNA from intact cells because: *i)* numerous studies have shown that the effects of PMA on intact microbial cells are minimal and PMA overwhelmingly targets extracellular DNA or DNA from dead cells (*15*, *16*, *18*); *ii)* our own tests confirmed PMA treatment did not lead to significant reductions in the amounts of DNA coming from live bacterial or fungal cells (Fig. S1B,C); and *iii)* if PMA inflated estimates of the amount of relic DNA by entering live cells, we would expect to detect relic DNA in all soils studied, which was not the case (Fig. 1A,B).

**Figure 1:**
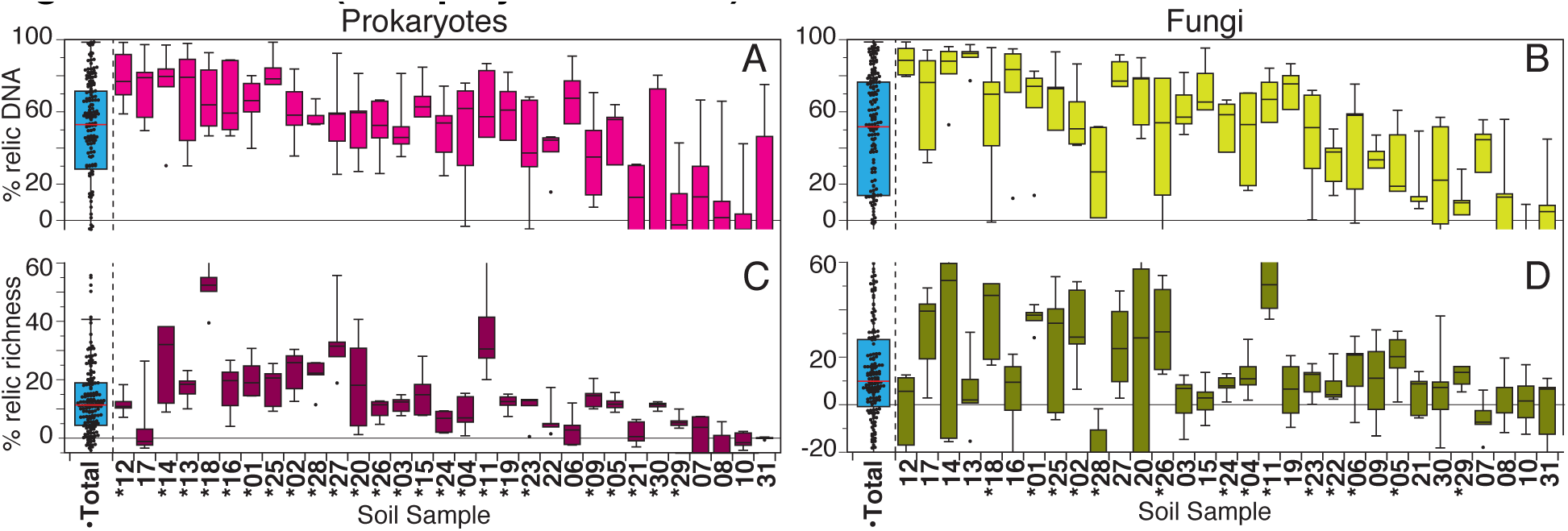
Relic DNA inflates estimation of soil microbial diversity. Percent of total prokaryotic 16S rRNA gene copies (A) or fungal ITS copies (B) in the relic DNA pool. Percent of total prokaryotic (C) or fungal (D) richness in relic DNA pool. Soils are ordered from left to right by decreasing mean percent prokaryotic relic DNA. (^*^) denotes significant differences in richness after relic DNA was removed (two-tailed itest *q* < 0.05). (^·^) Denotes significant richness differences after relic DNA removal across all soils (two-tailed itest *P* ≤ 0.05). Some box plots are truncated, see Supplementary Dataset S1 for complete dataset.

Removal of relic DNA significantly reduced estimates of microbial diversity. Across all samples, 13.9 ± 1.20% (mean ± SE; *n*=155) of the total prokaryotic richness (number of taxa) was found in the relic DNA pool (Fig.1C). In other words, nearly 14% of the taxa were no longer living in soil and were exclusively recovered in the relic DNA pool. In 24 of the soils tested, the prokaryotic richness was significantly lower, by as much as 55%, after relic DNA was removed (two-tailed *t* test *q* value ≤ 0.05) (Fig. 1C). The percent of prokaryotic taxa exclusively found in the relic DNA pool was positively correlated with the proportional abundance of 16S rRNA genes present in the relic DNA pool (Fig. S2A). That is, soils with more relic DNA tended to have lower richness once relic DNA was removed. Similar results were observed when we analyzed fungal DNA. The relic DNA pool contributed 12.4 ± 1.97% (mean ± SE; *n*=152) of the total fungal richness across all soils (Fig. 1D). The relic DNA contribution to estimates of fungal richness was significant in 14 soils (two-tailed *t* test *q* value ≤ 0.05), and removal of relic DNA reduced estimates of fungal diversity by up to 52% (Fig. 1D).

The removal of relic DNA can substantially reduce estimates of soil microbial diversity, indicating that the most commonly used molecular methods for assessing soil microbial diversity lead to inflated richness estimates due to the detection of DNA from dead cells. Although an average of ~40% of the total prokaryotic and fungal DNA was derived from relic DNA pools (Fig. 1A,B), the effect of relic DNA removal on fungal richness was variable across the soils examined (Fig. 1D) and there was no significant correlation between the percent of fungal relic DNA and the percent of fungal taxa found exclusively within the relic DNA pool (Fig. S2B). These between-sample differences in the magnitude of the effects of relic DNA on estimates of fungal diversity may be a product of differences in the temporal turnover of fungal communities at individual sites. If turnover in the fungal community composition is minimal, removal of relic DNA should have little effect on estimated fungal taxonomic richness as the relic DNA pool would reflect the diversity found in the pool of DNA extracted from intact fungal cells. This suggests that targeted analyses of the relic DNA pool could be used to identify taxa that once lived in soil, but are no longer alive due to changes in soil conditions, or to discriminate between endemic microbes and those microbes that have been transported into soils that can not support their growth.

We found that estimates of microbial community composition were also significantly influenced by the presence of relic DNA. Soil source was the strongest predictor of community differences (PERMANOVA R^2^=0.727, *P* ≤ 0.001 for prokaryotes; R^2^=0.646, *P* ≤ 0.001 for fungi). That is, we could discriminate between the distinct microbial communities found in the different soils, regardless of whether relic DNA was removed or not (Fig. S3A-D). However, the effect of relic DNA removal on community composition was significant for both prokaryotic and fungal communities (PERMANOVA R^2^=0.004, *P* ≤ 0.001 for prokaryotes; R^2^=0.002, *P* ≤ 0.001 for fungi). On an individual soil basis, the composition of prokaryotic communities was significantly affected by the removal of relic DNA in all 31 of the soils tested (PERMANOVA R^2^=0.10-0.23, *q* value ≤ 0.05) (Fig. 2A). In 21 of the 31 soils, removal of relic DNA also had a significant effect on the composition of fungal communities (PERMANOVA R^2^=0.10-0.22, *q* value ≤ 0.05) (Fig. 2B). The effects of relic DNA on the composition of both prokaryotic and fungal communities were positively correlated (Fig. 2C), highlighting that the magnitude of these relic DNA effects on microbial community composition were similar for both prokaryotic and fungal communities.

**Figure 2:**
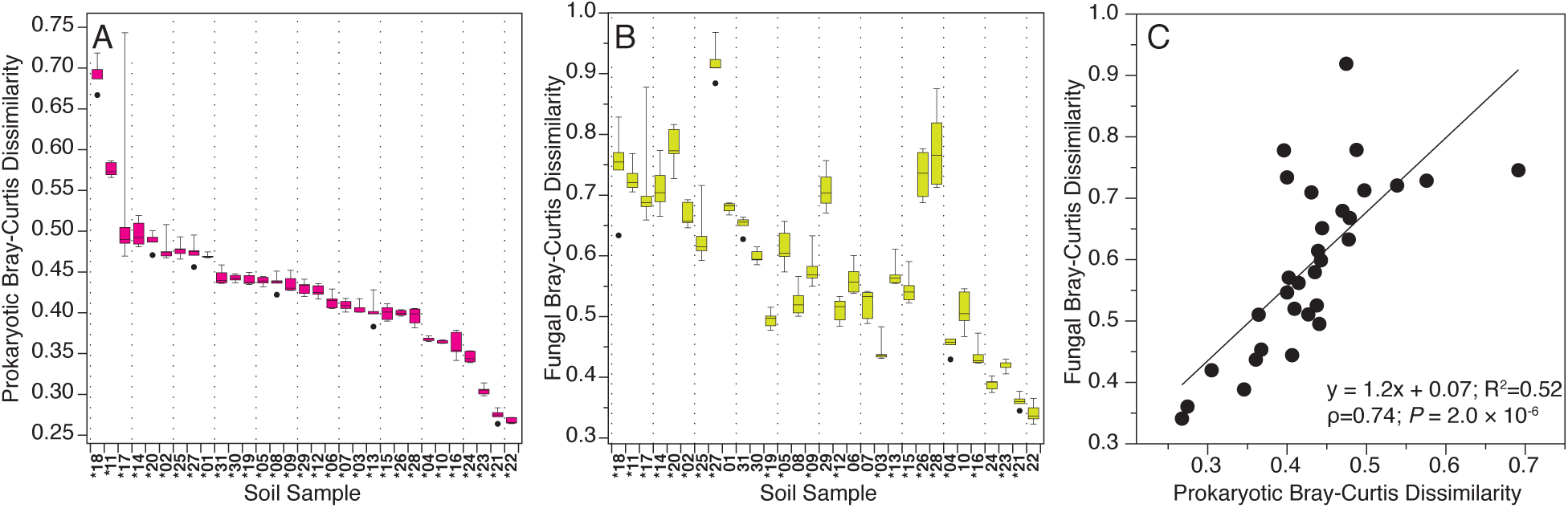
Relic DNA removal has a significant effect on community structure. Mean dissimilarity in soil prokaryotic (A) and fungal (B) communities after relic DNA removal, relative to untreated soils. Soils in A & B are ordered from left to right by decreasing order of the mean dissimilarity for prokaryotic communities. (^*^) denotes significant community differences between relic and total DNA pools (PERMANOVA *q* ≤ 0.05). C) Mean dissimilarity in prokaryotic communities is correlated with the associated dissimilarity in fungal communities.

The relative abundances of numerous key microbial lineages changed after the removal of relic DNA, but the taxa that changed, and the direction of observed shifts, varied across soils. For example, in a New Hampshire grassland (soil 25), Actinobacteria and α-Proteobacteria significantly increased in relative abundance after relic DNA was removed, but Verrucomicrobia decreased (Fig. 3). In many cases, the changes in estimated relative abundances after relic DNA removal were large, often approaching or exceeding 25% (Fig. 3). The relative abundances of α-Proteobacteria were consistently and significantly greater after relic DNA removal (Mann-Whitney *U* two-tailed *P* ≤ 0.05) (Fig. S4A), suggesting that the abundances of viable or dormant α-Proteobacteria are underestimated in many soil studies. In contrast, agaricomycete fungi were significantly less abundant after relic DNA was removed (Mann-Whitney *U* two-tailed *P* ≤ 0.05) (Fig. S4B), suggesting Agaricomycetes are less abundant than commonly thought. Together, these results show that the effects of relic DNA removal vary depending on the taxon in question, are not predictable *a priori*, and will vary depending on the particular soil studied.

**Figure 3:**
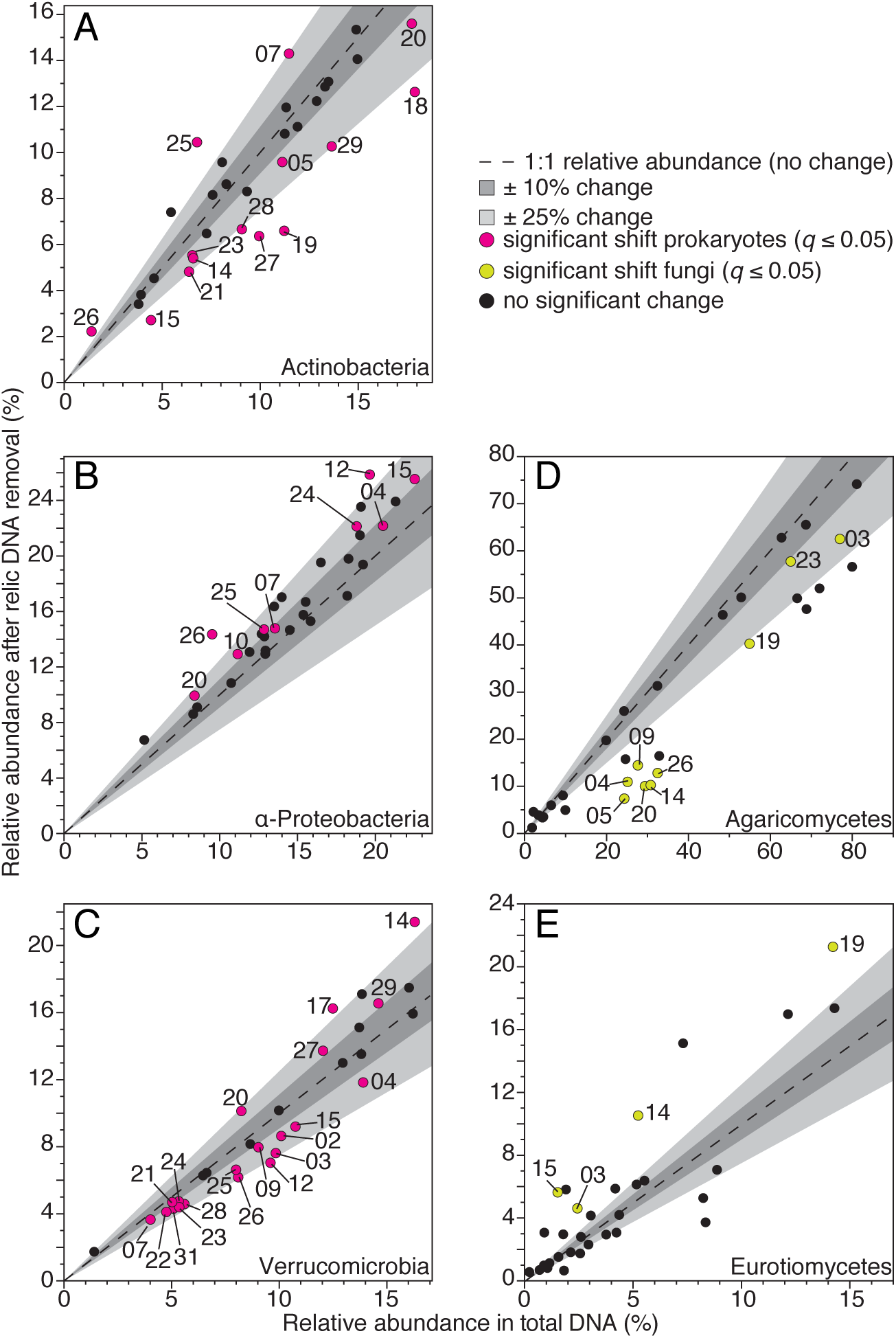
The relative abundances of key microbial lineages change after removal of relic DNA. Points are the mean relative abundances of Actinobacteria (A); α-Proteobacteria (B); Verrucomicrobia (C); Agaricomycetes (D); and Eurotiomycetes (E) before and after relic DNA removal. The relative abundances of colored points are significantly different (two-tailed Mann-Whitney U *q* ≤ 0.05) after relic DNA removal. Numbers correspond to soil sample (Supplementary Table 1). All significant changes in taxa comprising ≥ 5% of the total prokaryotic and ≥ 3% of the fungal communities across all soils are shown. In all plots, dashed lines represent no change in relative abundances (1:1 line). The dark grey shaded region represents ±10% change; light grey shaded region represents ±25% change. See Supplementary Dataset S1 for the mean relative abundances ±SE of each taxon in each soil.

Relating the abundances of microbial taxa or protein-coding genes to soil biogeochemical process rates has been challenging, hindering attempts to link microbial communities to the ecosystem-level processes they can control (*19*). Because the abundances of living microbial populations may be much higher or lower than is apparent from estimates of relative abundances derived from total DNA analyses, our results show that relic DNA likely obscures correlations between the abundances of individual microbial taxa (or their functional genes) and key biogeochemical processes. We found evidence of this when we compared the relative abundances of prokaryotes integral to soil nitrification before and after relic DNA was removed. For instance, ammonia oxidizing archaea classified as ‘*Ca*. Nitrosphaera’ and nitrite oxidizing bacteria classified as *Nitrospira spp.* changed by >25% in several soils after relic DNA was removed (Fig. S5A,B). Similarly, the relative abundance of *Glomeraceae*, a family of arbuscular mycorrhizal fungi, increased by >25% in 2 soils and decreased by >25% in 7 soils (Fig. S5C). Thus, by removing relic DNA prior to investigating relationships between specific soil processes and DNA-based quantification of microbial abundances, researchers may uncover more robust associations between microbes and key soil processes.

Because relic DNA can result in the overestimation of soil microbial diversity (Fig. 1), change assessments of overall community composition (Fig. 2), and alter the observed abundances of individual taxa (Fig. 3), we investigated which edaphic characteristics were predictive of the presence of relic DNA. Consistent with previous studies (reviewed in (*11*)), we show that edaphic characteristics influencing electrostatic interactions between DNA and soil particles were significant predictors of the presence of microbial relic DNA (Table 1). For example, soils with few exchangeable bases, especially K^+^ and Ca^2+^, were likely to contain relic DNA from both prokaryotic and fungal sources (Table 1, Fig S6). While soils with low pH, electrical conductivity and cation exchange capacity were more likely to harbor relic DNA, this pattern was stronger for prokaryotes than for fungi (Table 1). Moreover, pH predicted the change in both prokaryotic and fungal community composition after relic DNA removal (Fig. S7). These results highlight that, although the effects of relic DNA are variable across different soil types, it is especially important to account for relic DNA in acidic soils, or in soils with few exchangeable base cations (below *ca*. 40 meq 100 g^-1^).

**Table 1.** *P* values for logistic regression models fitting edaphic characteristics to the presence (≥ 20% of total DNA) or absence of relic DNA.

| Soil Characteristic | Prokaryotic | Fungal |
| --- | --- | --- |
| MWD<sub>w</sub> | 0.062 | N/S |
| pH | 0.048 | N/S |
| Electrical conductivity (mmhos cm<sup>-1</sup>) | 0.042 | N/S |
| NO<sub>3</sub><sup>-</sup>-N (ppm) | 0.014 | 0.025 |
| K (ppm) | 0.011 | 0.061 |
| Exchangeable Ca<sup>2+</sup> (meq 100 g<sup>-1</sup>) | 0.010 | 0.043 |
| Exchangeable Mg<sup>2+</sup> (meq 100 g<sup>-1</sup>) | 0.012 | N/S |
| Exchangeable Na<sup>+</sup> (meq 100 g<sup>-1</sup>) | 0.097 | 0.044 |
| Exchangeable K<sup>+</sup> (meq 100 g<sup>-1</sup>) | 0.008 | 0.022 |
| Total exchangeable bases (meq 100 g<sup>-1</sup>) | 0.008 | 0.036 |
| Cation exchange capacity (meq 100 g<sup>-1</sup>) | 0.046 | N/S |

All relationships are inverse, except Mean Weight Diameter (MWD<sub>w</sub>).
N/S: Not Significant at $P \leq 0.1$

Our finding that relic DNA can lead to significant overestimation of soil microbial diversity and reduce the ability to accurately quantify prokaryotic and fungal community structure has several important implications. First, it suggests that the actual microbial diversity in soil is lower than often reported. Second, relic DNA may obscure subtle spatiotemporal patterns or treatment effects in microbial communities. For example, shifts in soil microbial communities across seasons, or with plant species growing on a site, are often difficult to detect from DNA-based analyses (*20*, *21*). Similarly, long-term soil transplant experiments designed to study effects of climate change on soil microbial communities have showed little change in microbial community composition (*22*). Our ability to detect such shifts in soil microbial communities should increase if the ‘noise’ generated from non-living microbes is reduced by first removing relic DNA. Finally, the extreme diversity of soil microbial communities presents multiple computational problems for metagenomic assembly and analysis (*23*). Our data shows that a significant portion of this diversity is coming from relic DNA pools, suggesting that removal of relic DNA from samples prior to shotgun metagenomic analyses will result in improved metagenomic assemblies and improve our ability to infer the genomic attributes of undescribed soil microbes (*24*).

Relic DNA dynamics may have important ramifications for understanding community composition and processes in other ecosystems besides soil. For example, deep sea sediments also harbor large amounts of extracellular DNA (*25*), suggesting the removal of relic DNA would also affect diversity estimates in the deep biosphere. Specific analysis of the relic DNA pool in deep biosphere samples may have particular utility as a ‘fossil record’ to distinguish extinct microbial taxa from living organisms and more accurately reconstruct the subsurface paleome (*26*, *27*). More generally, relic DNA likely influences studies where DNA from dead organisms may be abundant or where DNA is particularly resistant to degradation, including studies of the microbial diversity found on aquatic particles, on mineral surfaces, in the human body, and in the built environment.

## Acknowledgements

Raw sequence data are available in the Sequence Read Archive at the National Center for Biotechnology Information (project accession no. SRP070563). We thank Stuart Grandy and Jörn Schnecker at University of New Hampshire, Cliff Bueno de Mesquita and Lara Vimercati at the University of Colorado Boulder, Eric Skokan at Black Cat Farms, and Kendra McLachlan at Kansas State University for soil collection or access to collection sites. We also thank Jessica Henley, Robin Hacker-Cary and Kristen Vaccarello for assistance with DNA extractions. Soil tests were preformed at Colorado State University’s Soil, Water and Plant Testing Laboratory. Sequencing was performed at the University of Colorado BioFrontiers Institute’s Next-Gen Sequencing Core Facility. Funding to support this work was provided by the National Science Foundation (DEB 0953331, EAR 1331828, DUE 1259336, EAR 1461281), and a Visiting Postdoctoral Fellowship award to P.C. from the Cooperative Institute for Research in Environmental Sciences.

## Supplementary Materials for

### Relic DNA is abundant in soil and obscures estimates of soil microbial diversity

Paul Carini^1^*, Patrick J. Marsden^2^, Jonathan W. Leff^1^,^3^, Emily E. Morgan^1^, Michael S. Strickland^4^, Noah Fierer^1^,^3^*

*Corresponding Authors: Paul Carini and Noah Fierer

Supporting Table 1: List of soil sample locations, descriptions and edaphic characteristics.

Supporting Dataset 1: Full dataset for Fig. 1 and mean % each taxa per soil, per treatment

### Materials and Methods

#### Soil collection and edaphic characteristics

Thirty one surface soils (0-5 cm; mineral soils only) were collected from locations in Colorado, New Hampshire, Virginia and Kansas, USA in August or September 2015 (Table S1), sieved to 2.0 mm, homogenized and stored at 4°C until PMA treatment. Soil characteristics were measured at the Colorado State University Soil Water and Plant Testing Laboratory using their standard protocols and included pH, electrical conductivity (mmhos cm^-1^), % organic matter, NO_3_- N (ppm), P (ppm), K (ppm), Zn (ppm), Fe (ppm), Mn (ppm), Cu (ppm), texture (% sand, silt, and clay), exchangeable bases (meq 100 g^-1^), and cation exchange capacity (meq 100 g^-1^). Percent moisture was determined gravimetrically on soils sieved to 2 mm before and after drying at 80°C for 72 hours. Mean weight diameter (MWD_W_; a proxy for the amount of water stable aggregates) was determined using the wet sieving method described in (*28*). Briefly, 2.0 g (W_t_) wet soils were sieved in a water bath through a 0.25 mm (d) sieve for 3 minutes. Aggregates remaining on the filter were dried at 80°C for 72 h and weighed (W_s_). Mean Weight Diameter after wet sieving (MWD_W_) = stable aggregates (W_s_)* sieve diameter (d) / Initial weight (W_t_).

#### Propidium monoazide treatment and DNA extraction

After homogenizing each soil, 10 replicate sub-samples of each soil type were resuspended (1% w/v) in sterile PBS (pH 7.4) in transparent screw cap tubes. PMA was added to 5 of the sub-samples (the PMA-treated samples) at a final concentration of 40 *µ*M in a dark room. The five PMA-treated and the five untreated sub-samples were vortexed (setting 6 on VWR-brand vortexer) for 4 minutes in the dark at room temperature. After the incubation, both PMA-treated and untreated samples were exposed to a 650-watt halogen lamp placed 20 cm from the sample tubes for four consecutive 30 s:30 s lightdark cycles, while continually vortexing to ensure even light exposure throughout resuspension. After light exposure, samples were frozen at ‐20°C until DNA extraction. DNA was extracted from 960 *µ*L of homogenized soil/PBS slurry using a MoBio PowerSoil DNA extraction kit, following manufacturer’s instructions. Experiments were conducted to determine the efficacy of this approach and specifically determine whether: i) there was excess capacity of 40 *µ*M PMA to remove naked DNA; and ii) if PMA penetrated living *Escherichia coli* or *Saccharomyces cerevisiae,* harvested in logarithmic-phase, and resuspended in PBS. Details and results of these control experiments are provided in Fig. S1.

#### Quantitative PCR

qPCR reactions were conducted in 25 μL total volumes on a Eppendorf realplex^2^ Mastercycler ep gradient S. The reaction mixture was as follows: 12.5 μl of Absolute qPCR SYBR Green Mix (Thermo Scientific) master mix; 1.25 μl of each primer (prokaryotic 16S rRNA: 515F 5’-GTGCCAGCMGCCGCGGTAA-3’ & 806R 5’-GGACTACHVGGGTWTCTAAT-3’; fungal ITS: 5’-CTTGGTCATTTAGAGGAAGTAA-3’ & ITS2 5’-GCTGCGTTCTTCATCGATGC-3’); 5 μl water; 5 μl of template DNA. All reactions were run in triplicate in 96 well plates with triplicate standard curves containing purified *E. coli* genomic DNA dilutions for 16S rRNA gene quantification and purified *Aspergillus fumigatus* (ATCC MYA-4609D-2) genomic DNA dilutions for ITS amplicon quantification. Program: 95°C for 15 min, followed by 40 cycles of (94°C 45 s; 50°C 60 s; 72°C 90 s) final extension 72°C 10 min. A qPCR ‘no template’ negative control was included with each qPCR run.

#### Amplicon sequencing and analytical methods

For sequence-based analyses of 16S rRNA and ITS gene regions (for prokaryotes and fungi respectively), we used the approaches described previously (*29*). Briefly, we amplified each of the 310 DNA samples in triplicate in 25 μL PCR reactions containing: 12.5 μl of Promega GoTaq Hot Start Colorless Master Mix; 0.5 μl of each barcoded primer (bacterial 16S: 515F 5’-GTGCCAGCMGCCGCGGTAA-3’ & 806R 5’-GGACTACHVGGGTWTCTAAT-3’; fungal ITS: 5’-CTTGGTCATTTAGAGGAAGTAA-3’ & ITS2 5’-GCTGCGTTCTTCATCGATGC-3’); 10.5 μl water; 1 μl of template DNA. Program: 94°C for 5 min, followed by 35 cycles of (94°C 45 s; 50°C 60 s; 72°C 90 s) and a final extension 72°C 10 min. Several negative controls, including ‘no template’ controls and ‘DNA extraction kit’ controls, were included alongside the soil DNA samples and sequenced. Amplicons were cleaned and normalized using the ThermoFisher Scientific SequalPrep Normalization Plate kit. Amplicons pools were spiked with phiX (15%) and sequenced on an Illumina MiSeq using v2 500-cycle Paired End kits at the University of Colorado BioFrontiers Institute’s Next-Gen Sequencing Core Facility. Reads were processed as described in (*29*). Briefly, raw amplicon sequences were demultiplexed according to the raw barcodes and processed with the UPARSE pipeline (*30*). A database of ≥97% similar sequence clusters was constructed in USEARCH (Version 8) (*31*) by merging paired end reads, using a “maxee” value of 0.5 when quality filtering sequences, dereplicating identical sequences, removing singleton sequences, clustering sequences after singleton removal, and filtering out cluster representative sequences that were not ≥75% similar to any sequence in Greengenes (for prokaryotes; Version 13_8) (*32*) or UNITE (for fungi) (*33*) databases. Demultiplexed sequences were mapped against the *de novo* constructed databases to generate counts of sequences matching clusters (i.e. taxa) for each sample. Taxonomy was assigned to each taxon using the RDP classifier with a threshold of 0.5 *(34)* and trained on the Greengages or UNITE databases. To normalize the sequencing depth across samples, samples were rarefied to 6,000 and 1,100 sequences per sample for the 16S rRNA and ITS analyses, respectively.

#### Statistical analyses

The percent changes in marker gene copy number and microbial richness in the relic DNA pool was calculated by comparing the mean number (n=5) of amplicons or taxa from total soil DNA extracts to the number of amplicons or taxa from each individual PMA treated subsample, using the formula (mean value in untreated samples - value in each PMA treated sample)/(mean value in untreated samples). Microbial richness and Bray-Curtis dissimilarities were calculated in the mctoolsr R package (*35*). Bray-Curtis dissimilarities were calculated on square root transformed taxa abundances. Where appropriate, statistical tests were corrected for multiple comparisons using the qvalue R package (*36*) using a significance threshold of *q* value ≤ 0.05. Spearman correlation coefficients and associated P values were calculated with the Hmisc R package (*37*). Logistic regression models were used to identify which edaphic characteristics predicted the presence of relic DNA (presence is defined as >20% relic DNA).

#### Rationale for using PMA vs alternative methods

Several techniques have been developed to investigate microbial viability in a variety of ecosystems (reviewed in (*17*)). The approach we used was selected because *i*) it can be used for high-throughput analyses; *ii)* the utility of PMA for the purposes of distinguishing extracellular DNA from DNA contained within cells is well documented and robust (*15*, *16*); and *iii)* key assumptions in other methods are untested. For example, we did not use RNA-based assays because it is impractical and prohibitively expensive to extract sufficient RNA from ~300 samples, as we do here. In addition, although RNA-based approaches are also thought to discriminate between metabolically active and inactive cells, this is not necessarily true (*38*). Moreover, the assumption that extracellular RNA cannot persist in soil has not been explicitly tested. It is possible that, as with DNA, there could be substantial amounts of RNA in soil coming from extracellular pools or cells that are no longer living, but additional studies are required to determine if this is the case.

**Figure S1:**
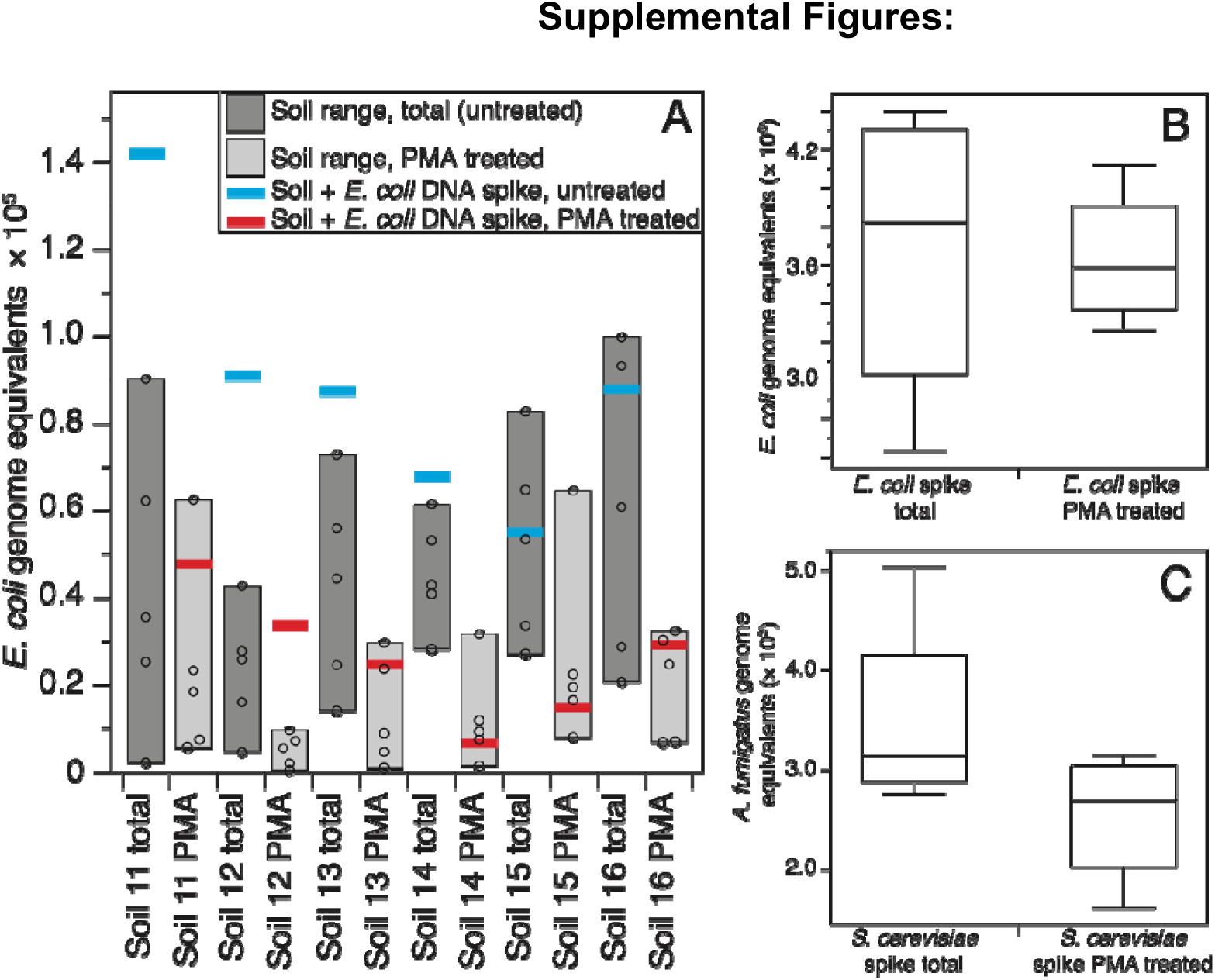
PMA effectively removed experimentally added naked DNA from some, but not all soils (A) and PMA did not penetrate live bacterial (B) or fungal (C) cells. A) Dark and light shaded boxes show the range of 16S rRNA gene copies detected from DNA extracts of untreated or PMA treated soils, respectively. Open circles are individual data points within range. Blue bars represent the number of 16S rRNA gene copies 4 quantified in fresh soil spiked with ~8.8 × 10^4^ copies of purified *E. coli* genomic DNA. Red bars represent the number of 16S rRNA gene copies quantified in soil spiked with 4 ~8.8 × 10^4^ copies of purified *E. coli* genomic DNA and treated with 40 *µ*M PMA as described for soils in ‘materials and methods’. PMA did not remove all of the spiked DNA in Soil 12, which also contained the largest percent of extracellular DNA (Fig. 1), suggesting additional PMA was necessary to remove the spiked DNA. B,C) Sterile PBS was spiked with equal volumes of exponentially growing *E. coli* and *S. cerevisiae* cell cultures. Tubes were either treated with 40 *µ*M PMA (n=5) or left untreated (n=5) as described for soils in ‘materials and methods’. The number of 16S rRNA gene copies or ITS amplicon copies were indistinguishable to untreated controls (two-tailed *t* test *P*=0.96 and *P*=0.22, respectively), illustrating PMA did not penetrate live bacterial or fungal cells.

**Figure S2:**
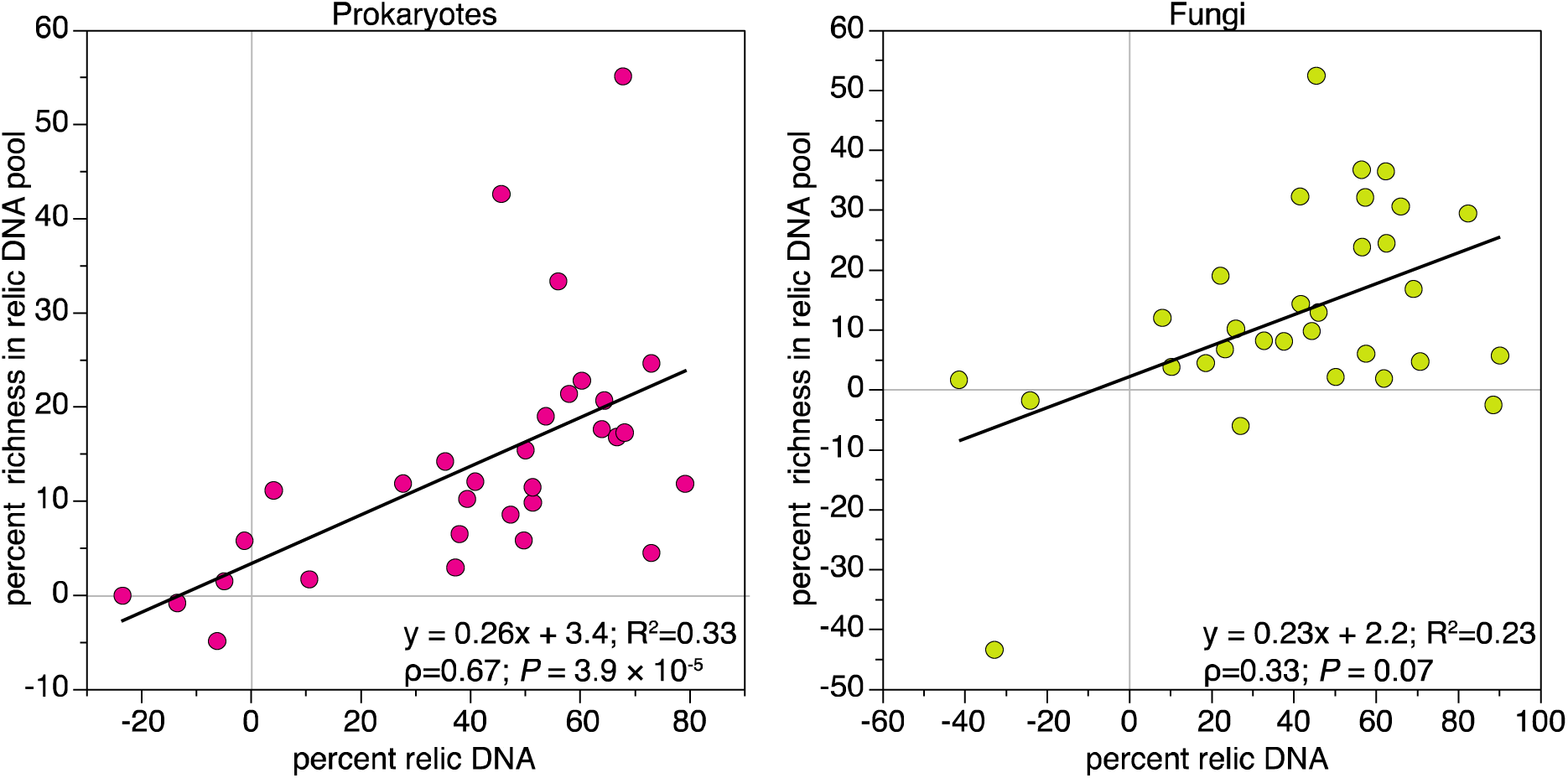
The percent of marker genes in the relic DNA pool is positively correlated with the proportional richness in the relic DNA pool, but this correlation is only significant for prokaryotes. Points are the mean percent relic DNA and mean percent richness in relic DNA, taken from Fig. 1. Linear regressions, formulas, Spearman’s *ρ* and *P* value are shown.

**Figure S3:**
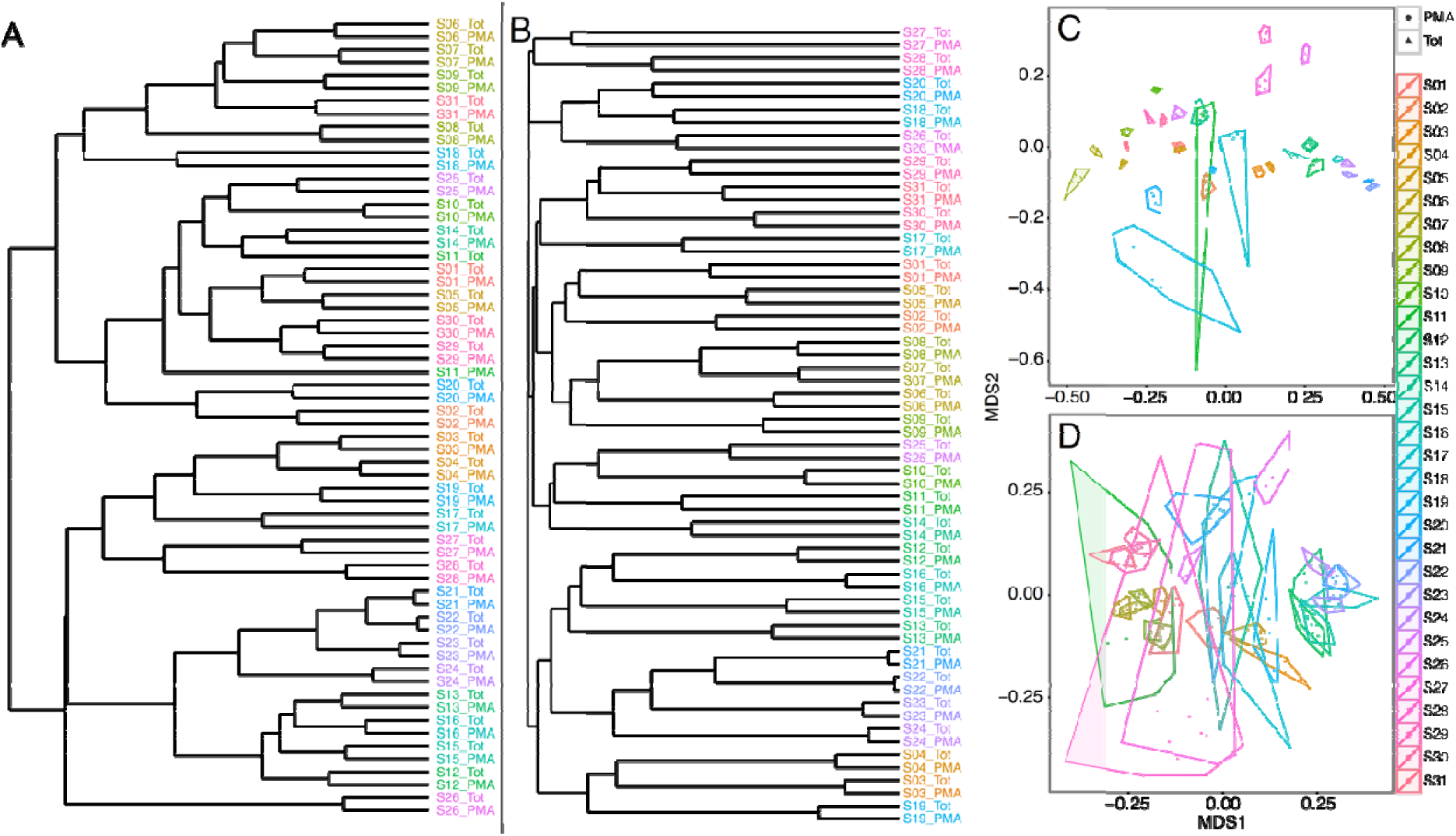
Differences in community composition between individual soil types is greater than the effect of relic DNA removal for any given soil. Dendrogram of prokaryotic (A) and fungal (B) community composition for all 31 soils. Multidimensional scaling plot of prokaryotic (C) and fungal (D) community composition for all 31 soils. Hulls in C & D connect the outermost points for each soil.

**Figure S4:**
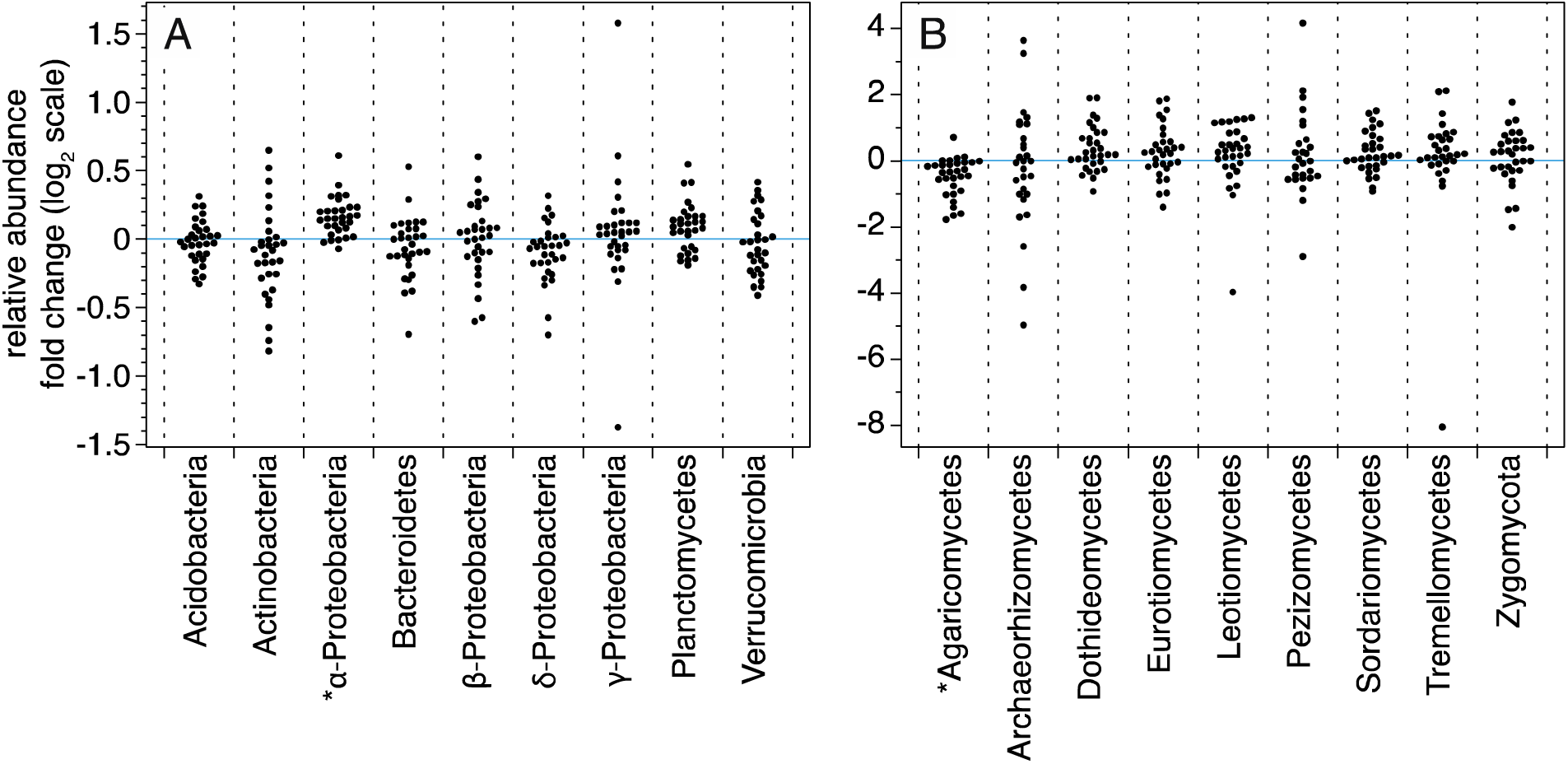
Fold changes in the estimated relative abundances of individual prokaryotic (A) and fungal (B) taxa after removal of relic DNA across all soils. Only major taxonomic groups comprising ≥5% of the total prokaryotic community or ≥3% of the total fungal community are shown. Points are the log_2_ fold change in estimated mean relative abundances after relic DNA removal. Points are not shown for some fungal taxa because they were not detected in one of the two conditions, preventing calculation of log_2_ fold change. See Supplemental Dataset S1 for full dataset. *Significant (two-tailed Mann-Whitney U *P*≤0.05).

**Figure S5:**
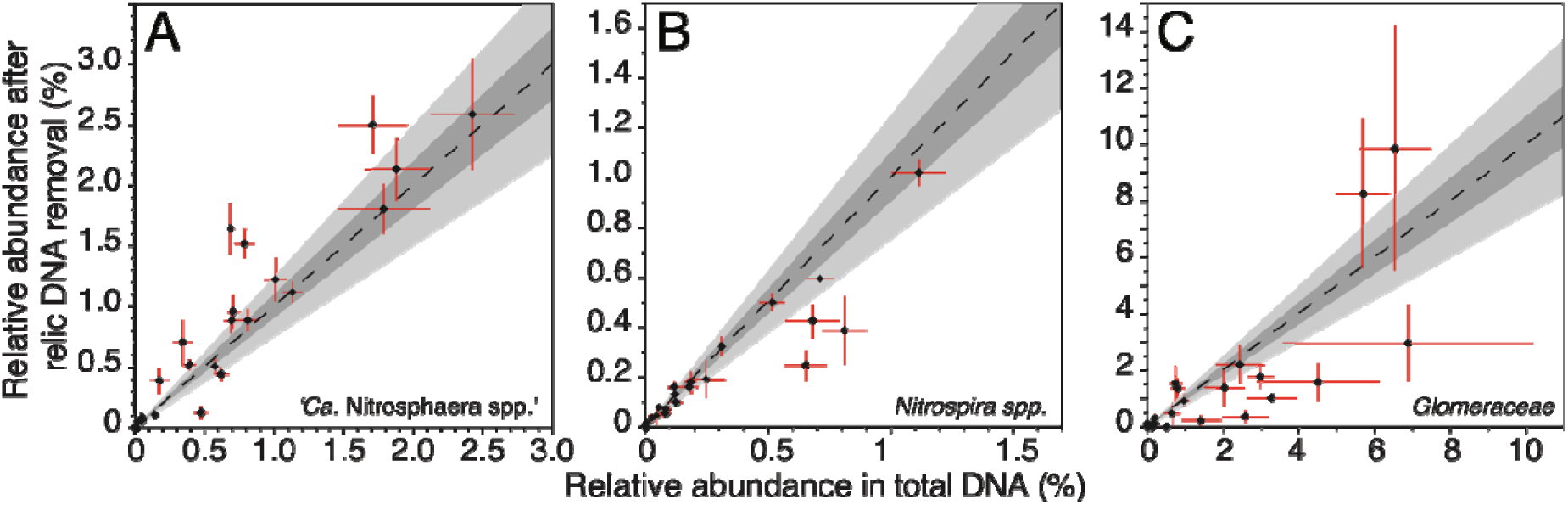
The relative abundances of prokaryotic nitrifiers and arbuscular mycorrhizal fungi change after relic DNA removal. Relative abundance of ammonia oxidizing archaea (A) or of nitrite oxidizing bacteria (B) before and after relic DNA removal (mean ± SE). C) Relative abundance of *Glomeraceae* (mycorrhizal fungi) before and after relic DNA removal (mean ± SE). In all plots, dashed line represents no change in relative abundance (1:1 line). Dark grey shaded region represents ±10% change; light grey shaded region represents ±25% change.

**Figure S6:**
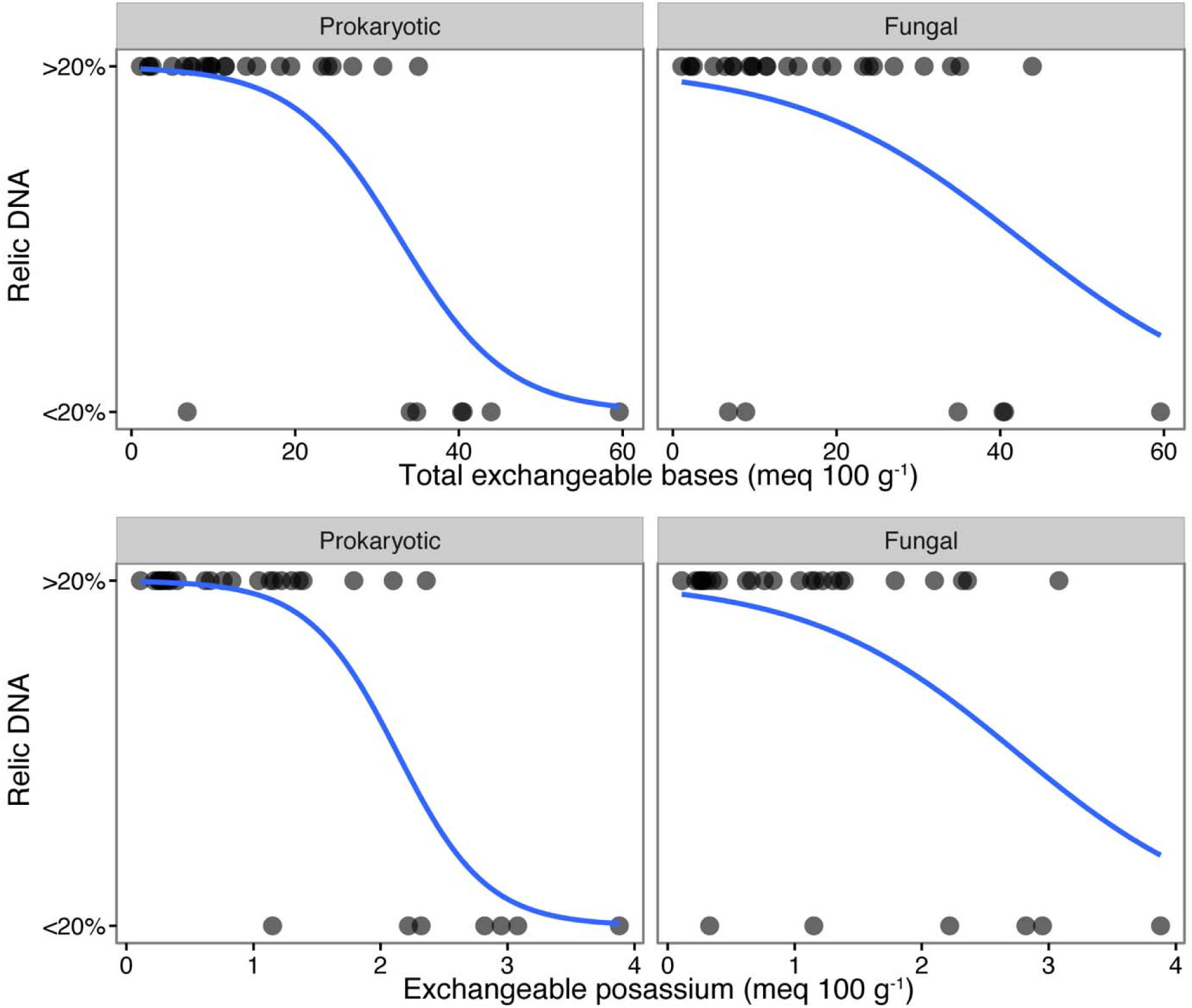
Logistic regressions fitting the presence of relic DNA to total exchangeable base cations (Na^+^, K^+^, Ca^2+^ and Mg^2+^) and exchangeable K^+^. Relic DNA was considered present if it represented ≥20% of the total DNA pool. Logistic regressions are shown in blue. Associated *P* values are provided in Table 1.

**Figure S7:**
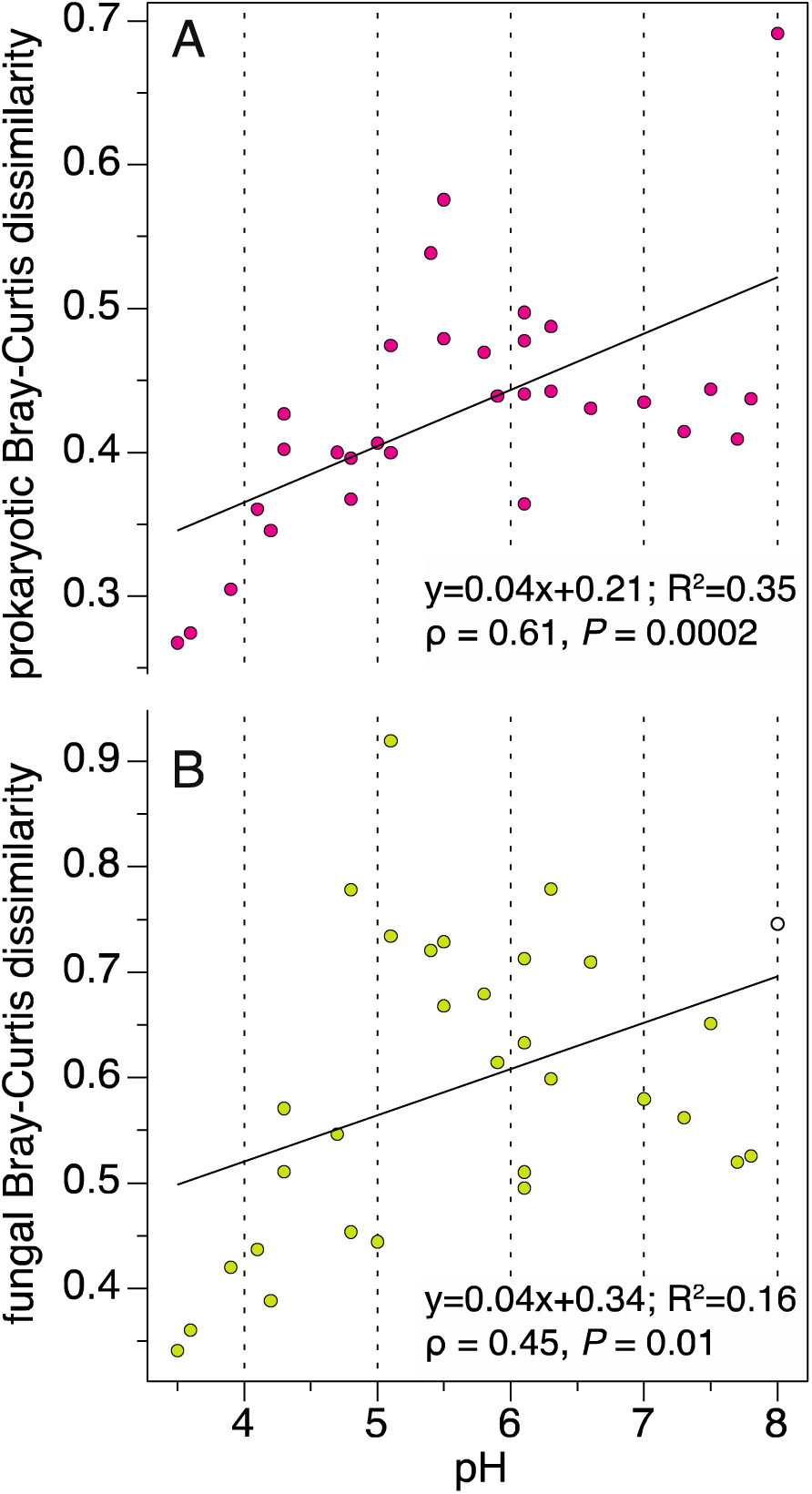
Soil pH is correlated with the difference in community composition after relic DNA removal. Points show the mean dissimilarity in soil microbial communities for prokaryotes (A) and fungi (B) after relic DNA removal, relative to total DNA extracts. Linear regressions, formulas, Spearman’s *ρ* and *P* values are shown.

